# High-throughput multiplexed tandem repeat genotyping using targeted long-read sequencing

**DOI:** 10.1101/673251

**Authors:** Devika Ganesamoorthy, Mengjia Yan, Valentine Murigneux, Chenxi Zhou, Minh Duc Cao, Tania PS Duarte, Lachlan JM Coin

**Affiliations:** Institute for Molecular Bioscience, The University of Queensland, Brisbane, QLD 4072, Australia

**Author notes:** Correspondence to Dr. Devika Ganesamoorthy, Institute for Molecular Bioscience, The University of Queensland, Brisbane, QLD 4072, Australia,; Prof. Lachlan JM Coin, Institute for Molecular Bioscience, The University of Queensland, Brisbane, QLD 4072, Australia.

**Keywords:** Tandem repeats, targeted sequencing, long-read sequencing

## Abstract

Tandem repeats (TRs) are highly prone to variation in copy numbers due to their repetitive and unstable nature, which makes them a major source of genomic variation between individuals. However, population variation of TRs have not been widely explored due to the limitations of existing tools, which are either low-throughput or restricted to a small subset of TRs. Here, we used SureSelect targeted sequencing approach combined with Nanopore sequencing to overcome these limitations. We achieved an average of 3062-fold target enrichment on a panel of 142 TR loci, generating an average of 97X sequence coverage on 7 samples utilizing 2 MinION flow-cells with 200ng of input DNA per sample. We identified a subset of 110 TR loci with length less than 2kb, and GC content greater than 25% for which we achieved an average genotyping rate of 75% and increasing to 91% for the highest-coverage sample. Alleles estimated from targeted long-read sequencing were concordant with gold standard PCR sizing analysis and moreover highly correlated with alleles estimated from whole genome long-read sequencing. We demonstrate a targeted long-read sequencing approach that enables simultaneous analysis of hundreds of TRs and accuracy is comparable to PCR sizing analysis. Our approach is feasible to scale for more targets and more samples facilitating large-scale analysis of TRs.

## INTRODUCTION

Repeated sequences occur in multiple copies throughout the genome, and they make up almost half of the human genome [1]. Repeat sequences can be divided into two categories, namely interspersed repeats and tandem repeats (TRs). Interspersed repeats are scattered throughout the genome and are remnants of transposons [2]. TRs consists of repeat units that are located adjacent to each other (*i.e.* in tandem). There are almost 1 million TRs in the human genome covering 10.6% of the entire genome [3]. TRs can be further divided into two types based on the length of the repeat unit; repeats with one to six base pair repeat units are classified as microsatellites or short tandem repeats (STRs) and those with more than six base pair repeat units are known as minisatellites [4].

TRs are prone to high rates of copy number variation and mutation due to the repetitive unstable nature, which makes them a major source of genomic variation between individuals. Variation in TRs may explain some of the phenotypic variation observed in complex diseases as it is poorly tagged by single nucleotide variation [5, 6]. Recent studies have shown that 10% to 20% of coding and regulatory regions contain TRs and suggested that variations in TRs could have phenotypic effect [7]. Although TRs represent a highly variable fraction of the genome, analysis of TRs so far are limited to known pathogenic regions, mainly STRs due to the limitations in analysis techniques.

Traditionally, TR analysis has been carried out via restriction fragment length polymorphism (RFLP) analysis [8] or PCR amplification of the target loci followed by fragment length analysis [9]. These techniques are only applicable to a specific target region and not scalable to high-throughput analysis, which limits the possibility of genome-wide TR analysis. In the recent decade, significant progress has been made in utilising high throughput short-read sequencing data for genotyping STRs [10]. Our group and others have also demonstrated targeted sequencing approaches using short-read sequencing for TR analysis [11, 12]. Several computational tools have been developed to improve the accuracy of TR genotyping from short-read sequencing data with varying performance [13–19]. Yet, most of these tools have focused mainly on the analysis of STRs and analysis of longer TRs remains a hurdle for these approaches. We reported GtTR in Ganesamoorthy *et. al.* (2018) [12], which utilizes short-read sequencing data to genotype longer TRs. GtTR reports absolute copy number of the TRs, but it does not report the exact genotype of two alleles due to the use of short-read sequencing data.

Sequencing reads that span the entire repeat region are informative to accurately genotype TRs [11], and therefore are ideal for genome-wide TR analysis. Long-read sequencing technologies have the potential to span all TRs in human genome, including long TRs. There have been few reports on the use of long-read sequencing for the analysis of specific TRs implicated in diseases [20–22]. Genotyping tools utilizing long-read sequencing data, such as Nanosatellite[21], RepeatHMM [23] and Tandem-genotypes [24] have been reported in the recent years with varying performance across different length of repeat units and repeat length. We reported VNTRTyper in Ganesamoorthy *et. al.* (2018) [12] to genotype TRs from long-read whole genome sequencing data. Despite the availability of genotyping tools, long-read sequencing is not widely used for TR analysis, due to the high costs associated with whole genome long-read sequencing. Cost-effective long-read sequencing approaches will be an important and attractive option to genotype TRs in large-scale studies. However, there has been limited progression on targeted long-read sequencing of TRs.

We have previously demonstrated that targeted sequence capture of repetitive TR sequences are feasible using short-read sequencing technologies [12]. In this study, we demonstrate the targeted sequence capture of repetitive TRs using Oxford Nanopore long-read sequencing technologies. There have been previous reports on the use of targeted sequencing combined with long read sequencing technologies [25], however enrichment of repeat sequences requires optimization in probe design and probe hybridization approaches. We optimized the protocols and report successful enrichment of repetitive sequences followed by long-read sequencing. We demonstrate the accuracy of genotype estimates from targeted long-read sequencing by comparison with gold-standard PCR sizing analysis. In this study, we predominantly targeted longer TRs (i.e. minisatellites), however our approach is applicable to all TRs. Our targeted long-read sequencing method presented here provides an accurate and cost effective approach for large-scale analysis of TRs, which will be useful for researchers to explore the impact of TR variants on diseases and phenotypes.

## RESULTS

We demonstrate a targeted sequence capture approach combined with Nanopore long-read sequencing to genotype hundreds of TRs.

### Targeted Capture Sequencing of Tandem repeats

We performed sequence enrichment of targeted TRs for 7 samples followed by long-read sequencing using Oxford Nanopore Technologies’ MinION as described in Methods. Figure 1a shows the read length distribution observed in targeted capture sequencing data. The median read length followed the expected read-length distribution, with the exception of an under-representation of repeats of length >3kb (Figure 1a). The read length in this study was sufficient to analyse majority of targeted TRs of length less than 2kb. Sequence coverage varied across targets and samples, on average 97X sequence coverage was achieved, with only 19 targets having less than 1X coverage (Figure 1b) and majority of the low coverage targets (16 of the 19 targets) have less than 25% GC content (Supplementary Figure 1)

**Figure 1:**
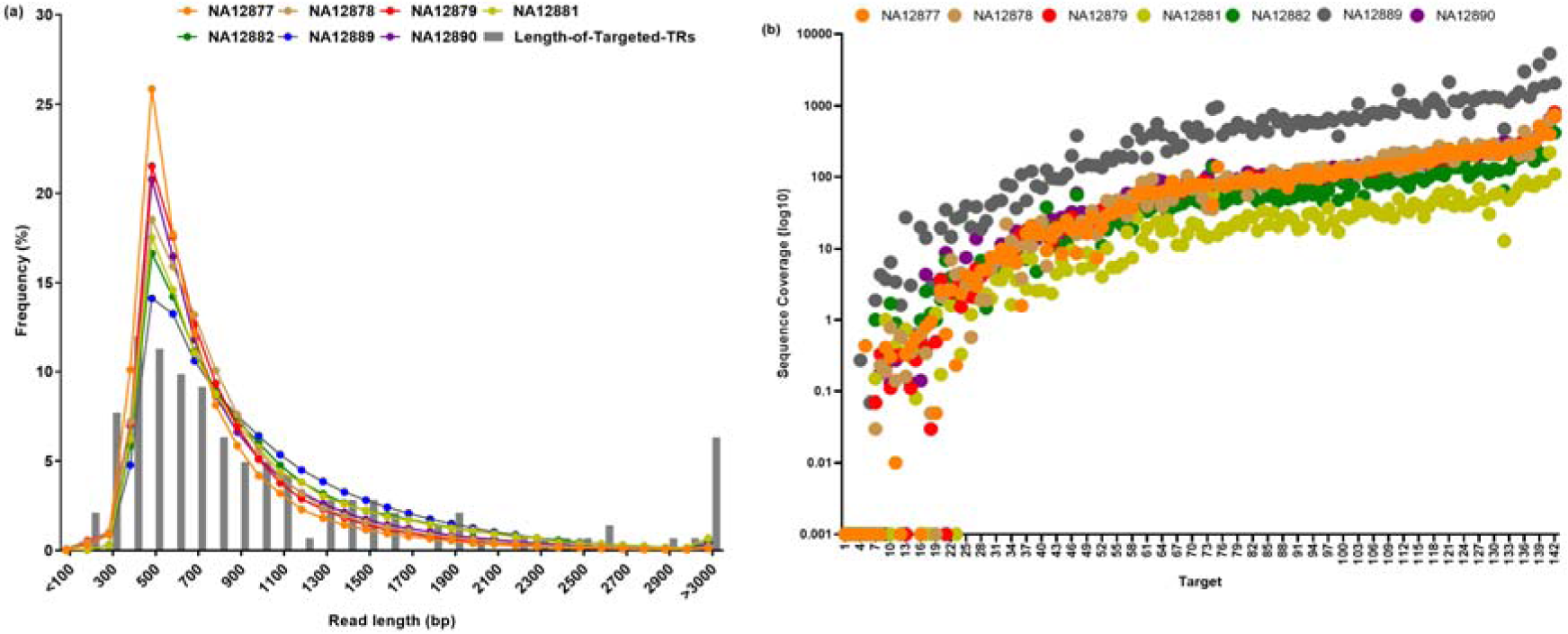
Read length and sequence coverage distribution (a) Read length distribution of Nanopore targeted sequencing. Lines indicates the read length distribtuion for each sample and grey bars indicate the length distribution of targeted TRs and (b) Sequence coverage distribution of Nanopore targeted sequencing for all 7 samples.

Supplementary Table 1 summarises the metrics for targeted sequencing on Nanopore sequencing technologies. Nanopore multiplexing (See Methods) group 1 samples had similar yield between samples, however, Nanopore multiplexing group 2 samples had varying yield per sample. Despite the differences in sequencing yield, we achieved an average of 3062-fold target enrichment and on target capture rate was approximately 50%.

### Genotyping of Tandem repeats using targeted long-read sequencing

Genotype estimates from targeted long-read sequencing datasets were estimated using our tool VNTRTyper [12] with the improvements described in Methods. We also applied Tandem-genotypes [24] to determine the genotypes of TRs from long-read sequencing data. We used a minimum of two reads as read threshold to determine the repeat number for each alleles.

Prior to obtaining any sequence data, we generated PCR sizing results as a gold standard on 10 targets for comparison to sequencing analysis. These 10 targets were selected to include various repeat unit length and repeat sequence combinations to assess the accuracy of the genotypes determined from sequencing data. Of these 10 targets, 2 were excluded for comparison as all 7 samples had insufficient number of spanning reads (minimum of 2 reads required for genotyping) to genotype these targets. Genotype estimates from VNTRtyper on these 8 targets correlated well with PCR (Pearson correlation greater than 0.980 for all samples) (Table 1 and Supplementary Figure 2). Genotype estimates by Tandem-genotypes also correlated well with PCR with correlation greater than 0.984 for all samples (Supplementary Table 2 and Supplementary Figure 3), however fewer targets had sufficient data to compare with PCR sizing results.

**Table 1:** Genotype estimates on Nanopore targeted capture sequencing using VNTRTyper

| Sample | Method | Genotype of Target* |  |  |  |  |  |  |  | Pearson<br>Correlation<br>with PCR |
| --- | --- | --- | --- | --- | --- | --- | --- | --- | --- | --- |
|  |  | TR_8<br>(12.0) | TR_57<br>(15.6) | TR_86<br>(2.0) | TR_87<br>(9.0) | TR_93<br>(2.0) | TR_109<br>(15.3) | TR_112<br>(4.0) | TR_120<br>(2.2) |  |
| NA12877 | PCR | 12.0/13.0 | 10.6/13.6 | 2.0/2.0 | 6.0/8.0 | 2.0/2.0 | 15.3/17.3 | 3.0/4.0 | 2.2/2.2 | 0.9988 |
|  | Nanopore | 12.0/13.0 | ND | 2.0/2.0 | 6.2/8.4 | 2.0/2.0 | 15.3/17.3 | 3.0/4.1 | 2.2/3.2 |  |
| NA12878 | PCR | 12.0/12.0 | 10.6/12.6 | 2.0/2.0 | 6.0/9.0 | 2.0/2.0 | 15.3/17.3 | 3.0/4.0 | 2.2/2.2 | 0.9978 |
|  | Nanopore | 11.0/12.1 | ND | 2.0/2.0 | 6.0/8.8 | 2.0/2.0 | 14.8 /17.1 | 3.0/4.0 | 2.2/3.2 |  |
| NA12879 | PCR | 12.0/13.0 | 10.6/10.6 | 2.0/2.0 | 8.0/9.0 | 2.0/2.0 | 15.3/17.3 | 3.0/3.0 | 2.2/2.2 | 0.9927 |
|  | Nanopore | 12.4/12.4 | 10.6/10.6 | 2.0/2.0 | 6.0/8.6 | 2.0/2.0 | 14.8/16.9 | 3.0/4.0 | 2.2/3.2 |  |
| NA12881 | PCR | 12.0/12.0 | 10.6/10.6 | 2.0/2.0 | 8.0/9.0 | 2.0/2.0 | 15.3/15.3 | 3.0/4.0 | 2.2/2.2 | 0.9944 |
|  | Nanopore | 12.0/12.0 | 10.6/10.6 | 2.0/2.0 | 8.0/9.0 | ND | 15.3/17.3 | 3.0/4.0 | 2.2/3.2 |  |
| NA12882 | PCR | 12.0/13.0 | 10.6/10.6 | 2.0/2.0 | 6.0/6.0 | 2.0/2.0 | 15.3/17.3 | 3.0/3.0 | 2.2/2.2 | 0.9919 |
|  | Nanopore | 11.8/13.0 | 10.6/10.6 | 2.0/2.0 | 6.0/8.5 | 2.0/2.0 | 15.3/17.3 | 3.0/4.0 | 2.2/3.2 |  |
| NA12889 | PCR | 12.0/12.0 | 13.6/17.6 | 2.0/2.0 | 8.0/8.0 | 2.0/2.0 | 17.3/17.3 | 4.0/4.0 | 2.2/2.2 | 0.9800 |
|  | Nanopore | 12.1/12.1 | 10.6/15.8 | 2.0/2.0 | 6.1/8.1 | 2.0/2.0 | 14.8/17.2 | 3.0/4.0 | 2.2/3.2 |  |
| NA12890 | PCR | 12.0/13.0 | 10.6/10.6 | 2.0/2.0 | 6.0/9.0 | 2.0/2.0 | 15.3/15.3 | 3.0/3.0 | 2.2/2.2 | 0.9936 |
|  | Nanopore | 12.2/12.2 | 10.6/10.6 | 2.0/2.0 | 6.0/9.0 | 2.0/2.0 | 13.3/15.3 | 3.0/4.0 | 2.2/3.2 |  |

\*Repeat Number in reference hg19 is provided within brackets for each target
\*Repeat numbers that do not agree with PCR results are highlighted in red.
ND – Sufficient data not available for genotype analysis

Genotype estimates from VNTRTyper and Tandem-genotypes for all 142 targets from Nanopore capture sequencing samples are provided in Supplementary Spreadsheets Table 1 and Table 2 respectively. Genotype estimates by VNTRtyper and Tandem-genotypes correlate well and the correlation values range from 0.904 to 0.994 for Nanopore targeted sequencing.

We were able to determine the genotype on average for 60% of the targets (range 48% to 75%) using VNTRTyper and 57% of the targets (range 41% to 74%) using Tandem-genotypes. Both VNTRTyper and Tandem-genotypes failed to genotype targets with low GC sequence content (< 25% GC content) and targets which are greater than 2Kb in length, which accounts for approximately 22% of the targets (32 of the 142 targets). Targets with low GC sequence content (< 25% GC content) didn’t have sufficient sequence coverage for analysis due to inefficient sequence enrichment in these regions (Supplementary Figure 1). Targets which are greater than 2Kb in length didn’t have sufficient spanning reads for genotyping analysis (See Methods and Figure 1a).

It was evident that the GC content of the target and size (i.e. repeat length) affected the genotyping efficiency of our targeted capture sequencing approach. Therefore, we assessed the genotyping rate based on the size of the target and GC content of the target (Figure 2). For all 142 targets genotyping rate using VNTRTyper was only 59.8% (Figure 2a), however genotyping rate improved to 67% for 125 targets with a size threshold of 2Kb (Figure 2b) and 67.1% for 125 targets with 25% GC threshold (Figure 2c). Furthermore, genotyping rate improved to 75.2% for 110 targets with a combined 2Kb size threshold and 25% GC threshold (Figure 2d). Also, sample with high sequence coverage (NA12889) had the highest genotyping rate of 90.9% for 110 targets (Supplementary Figure 4). Genotyping rate using Tandem-genotypes also improved to an average of 63.7% for 110 targets (range 43.6% to 85.5%) (Supplementary Figure 5).

**Figure 2:**
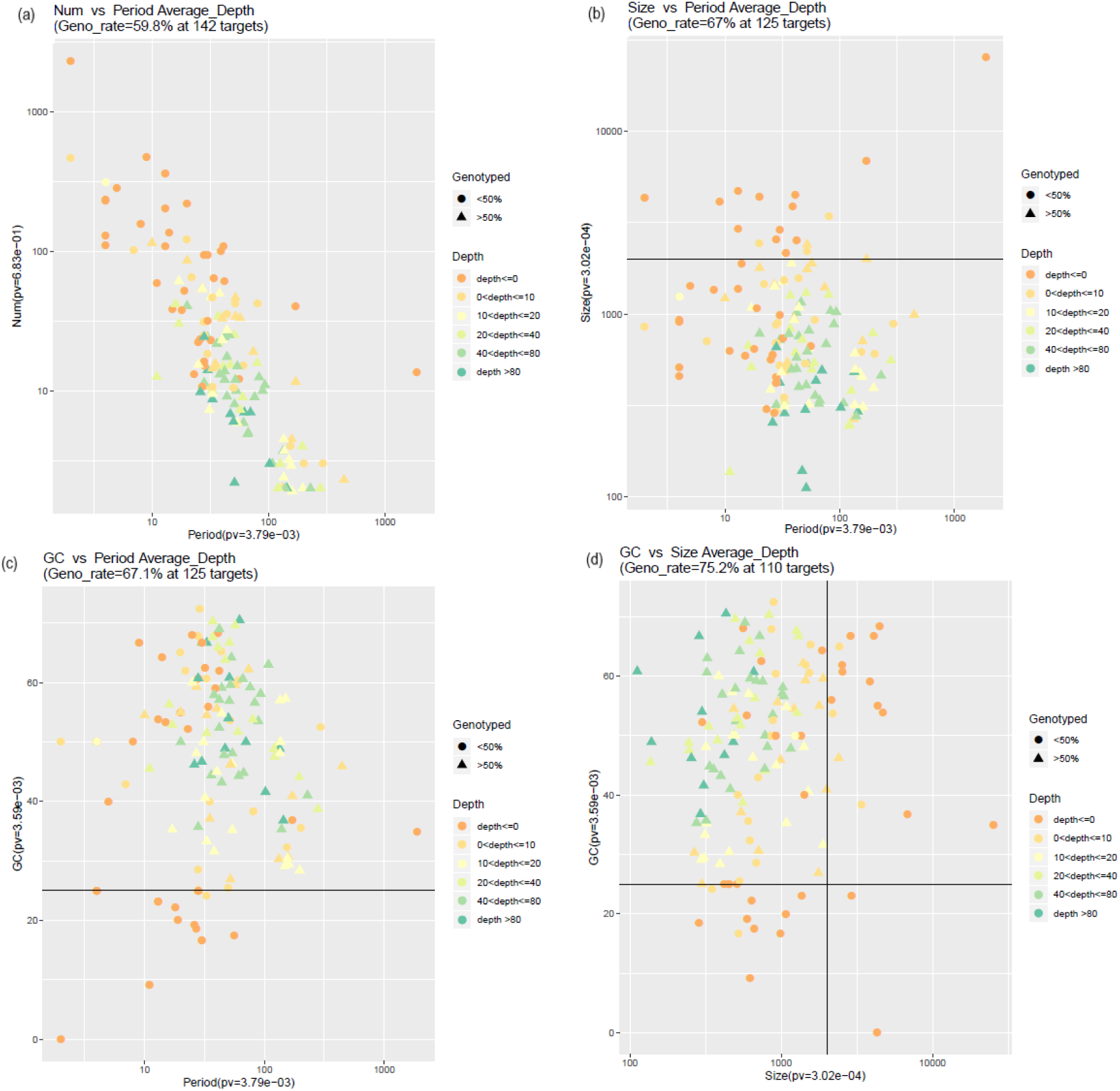
Assessment of genotyping rate using VNTRTyper based on the size and GC content of the target for all 7 samples. Triangle indicates that greater than 50% of the samples had a genotyping estimate and circle indicates only less than 50% of the samples had a genotyping estimate for the given target. Colours indicate the depth, which is defined as the number of spanning reads detected for the target region. Black thick lines inside the plots indicate the 2Kb size threshold and 25% GC content size threshold. (a) genotyping rate for all targets, shown as repeat number vs period (i.e. repeat unit), (b) genotyping rate with 2Kb size threshold, (c) genotyping rate with 25% GC threshold and (d) genotyping rate with both 25% GC threshold and 2Kb size threshold.

### Genotyping of Tandem repeats using long-read whole genome sequencing

To investigate the accuracy of genotype estimates of TRs from targeted sequence capture compared to WGS, we performed genotyping analysis on the targeted regions using VNTRTyper and Tandem-genotypes on whole genome long-read sequencing data. We downloaded whole genome long-read Nanopore and PacBio sequencing data on CEPH Pedigree 1463 NA12878 sample. We have previously reported genotyping estimates by VNTRTyper on PacBio NA12878 WGS data [12]. Here we use the genotype estimates by VNTRTyper on PacBio NA12878 WGS data to compare genotype estimates by Tandem-genotype and the results of targeted sequencing analysis.

We compared the accuracy of genotype estimates from WGS data with PCR sizing analysis. Genotype estimates by VNTRTyper and Tandem-genotypes on WGS data were compared with PCR sizing results on 10 targets (Table 2). VNTRTyper and Tandem-genotypes had comparable correlation with PCR sizing analysis for both Nanopore and PacBio WGS. Genotype estimates for all 142 targets from Nanopore and PacBio WGS data determined using VNTRTyper and Tandem-genotypes are provided in Supplementary Spreadsheets Table 3.

**Table 2:** Genotype estimates on NA12878 WGS and Capture sequencing data using VNTRTyper and Tandem-genotypes

| Method^ | Genotype of Target* |  |  |  |  |  |  |  |  |  | Pearson |
| --- | --- | --- | --- | --- | --- | --- | --- | --- | --- | --- | --- |
|  | TR_8 | TR_32 | TR_57 | TR_64 | TR_86 | TR_87 | TR_93 | TR_109 | TR_112 | TR_120 | Correlation<br>with PCR |
|  | (12.0) | (38.4) | (15.6) | (18.3) | (2.0) | (9.0) | (2.0) | (15.3) | (6.0) | (2.2) |  |
| PCR | 12.0/12.0 | 9.4/10.4 | 10.6/12.6 | 17.3/18.3 | 2.0/2.0 | 6.0/9.0 | 2.0/2.0 | 15.3/17.3 | 3.0/4.0 | 2.2/2.2 |  |
| NPW_VT | 12.0/12.0 | 8.2/10.7 | 10.6/12.6 | 17.3/18.6 | 2.0/2.0 | 6.0/9.0 | 2.0/2.0 | 15.3/17.3 | 3.0/4.0 | 2.2/3.2 | 0.9980 |
| NPW_TG | 11.0/12.0 | 4.4/9.4 | 10.6/12.6 | 18.3/19.3 | 2.0/2.0 | 6.0/9.0 | 2.0/2.0 | 15.3/17.3 | 3.0/4.0 | 2.2/2.2 | 0.9800 |
| PBW_VT | 12.0/12.0 | 11.0/12.4 | 10.6/12.6 | 18.3/19.3 | 2.0/2.0 | 6.0/9.0 | 2.0/2.0 | 15.3/17.3 | 3.0/4.0 | 2.2/2.2 | 0.9957 |
| PBW_TG | 12.0/12.0 | ND | 12.6/12.6 | ND | 2.0/2.0 | ND | 2.0/2.0 | 15.3/15.3 | 4.0/4.0 | 2.2/2.2 | 0.9900 |
| NPC_VT | 11.0/12.1 | ND | ND | ND | 2.0/2.0 | 6.0/8.8 | 2.0/2.0 | 14.8/17.1 | 3.0/4.0 | 2.2/3.2 | 0.9978 |
| NPC_TG | 11.0/12.0 | ND | ND | ND | 2.0/2.0 | 6.0/9.0 | ND | 15.3/17.3 | 3.0/4.0 | 2.2/2.2 | 0.9988 |

^NPW\_VT – Nanopore WGS VNTRTyper; NPW\_TG – Nanopore WGS Tandem-genotypes; PBW\_VT – PacBio WGS VNTRTyper; PBW\_TG – PacBio WGS Tandem-genotypes; NPC\_VT – Nanopore Capture sequencing VNTRTyper; NPC\_TG – Nanopore Capture sequencing Tandem-genotypes
\*Repeat Number in reference hg19 is provided within brackets for each target
\*Repeat numbers that do not agree with PCR results are highlighted in red.
ND – Sufficient data not available for genotype analysis

We compared the genotype estimates between WGS data and targeted capture sequencing data (77 targets which had results for both WGS and targeted sequencing). Genotype estimates by VNTRTyper between WGS data and targeted capture sequencing data showed a correlation of 0.9782 (correlation on 154 alleles) (Figure 3a). Genotype estimates by Tandem-genotypes had lower correlation between WGS and targeted capture sequencing data of 0.7694 (correlation on 152 alleles – 76 targets) (Figure 3b). On the subset of 7 targets for which we had generated PCR sizing analysis, Nanopore WGS data correlated with 12/14 genotype estimates on Nanopore capture sequencing using VNTRTyper precisely compared to PCR sizing (Table 2 and Figure 4a). Genotype estimates using Tandem-genotypes on Nanopore WGS data correlated with 11/12 genotype estimates on Nanopore capture sequencing precisely compared to PCR sizing (Figure 4b).

**Figure 3:**
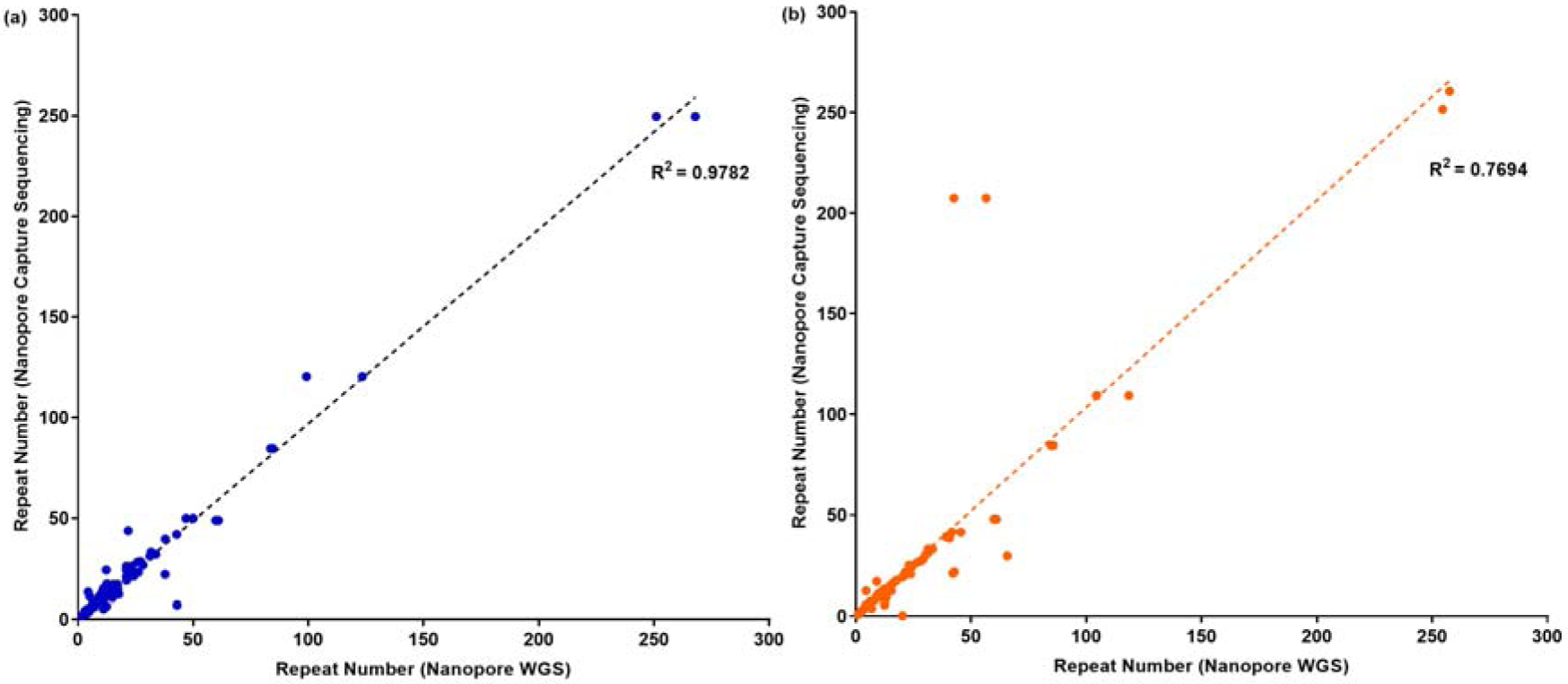
Correlation between WGS and Targeted sequencing genotype estimates using (a) VNTRTyper and (b) Tandem-genotypes for NA12878 sample.

**Figure 4:**
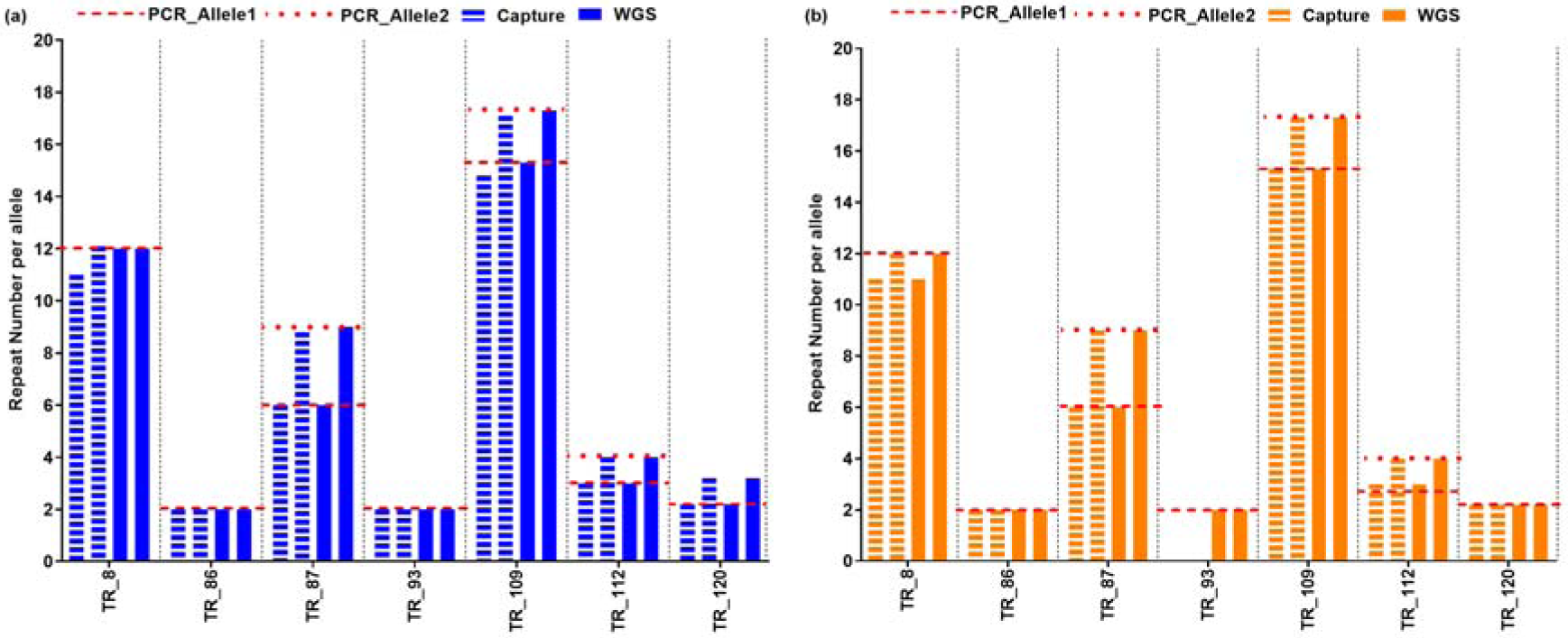
Comparison of genotype estimates between WGS and Target capture sequencing for NA12878 sample using (a) VNTRTyper and (b) Tandem-genotypes. Red line indicates PCR sizing results. Targets with no genotype estimates are shown as a gap for the corresponding column.

### Variation in Tandem repeats

To assess the extent of variation in repeat numbers between individuals, we compared the genotype estimates to the reported reference (hg19) repeat number. Genotype estimates determined by VNTRTyper on Nanopore capture sequencing on 7 members of CEPH pedigree 1463 were used to assess the variation. We found that for a given sample, on average 51% (range 45% - 60%) of the targets have a genotype which is different to the reference, with more deletions (28%) than duplications (23%) (Figure 5).

**Figure 5:**
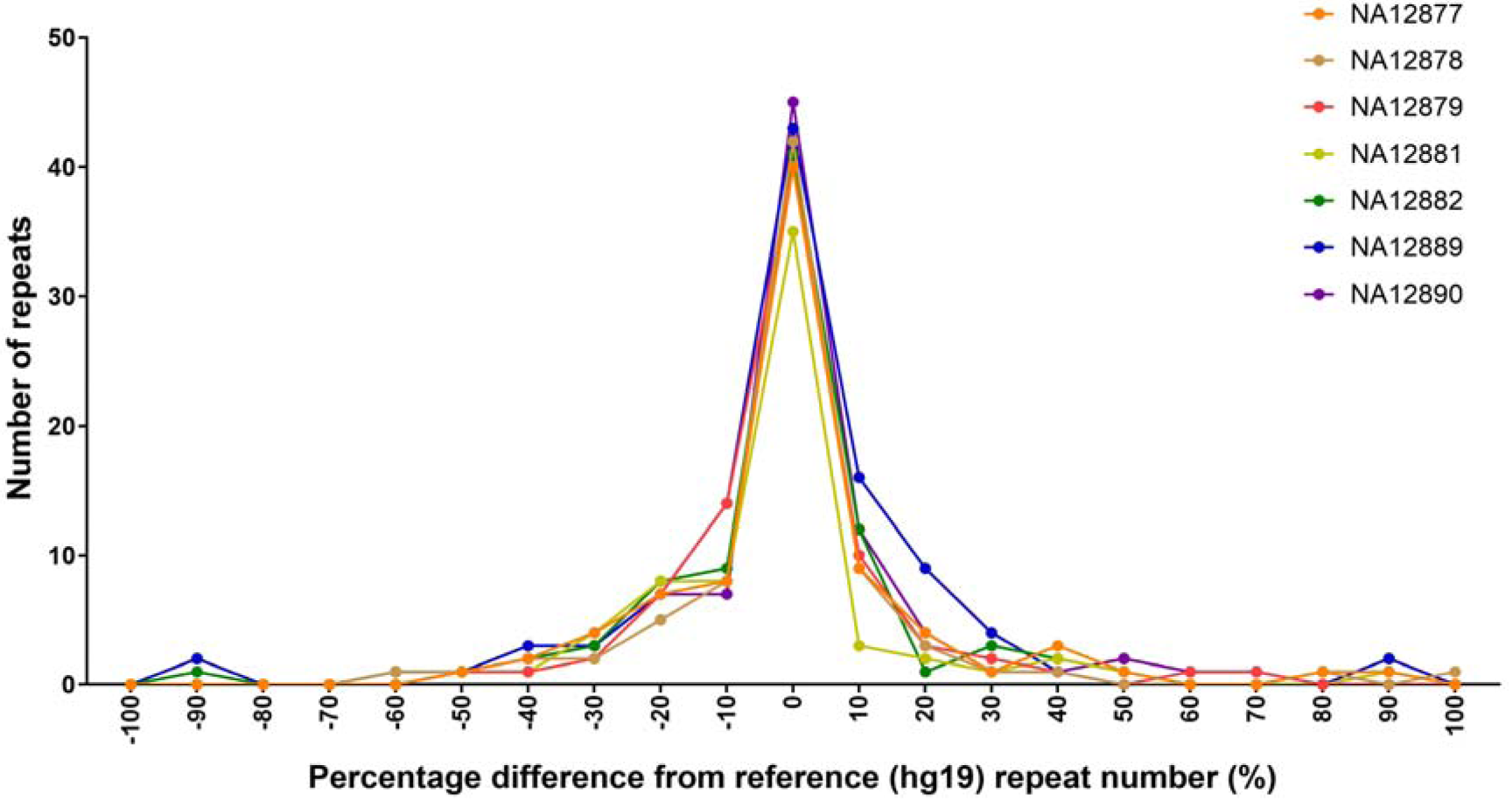
Percentage difference between reported repeat number in reference genome (hg19) and estimated repeat number based on genotype estimates using VNTRTyper on Nanopore targeted sequencing.

## DISCUSSION

In this study, we presented a targeted sequencing approach combined with long-read sequencing technology to genotype TRs. To our knowledge, this is the first report on genotyping analysis of hundreds of TRs using targeted long-read sequencing approach. Sequencing reads that span the entire repeat region and flanking region are often useful in providing an accurate estimation of the repeat genotype. Long-read sequencing technologies have the ability to generate reads which can span the entire repeat region and flanking regions. However, whole genome long-read sequencing analysis is still expensive for large-scale population analysis; hence, we developed a targeted long-read sequencing approach for TR analysis.

We showed that 1) target enrichment of repetitive sequences followed by long-read sequencing is feasible and 2) genotype predictions on targeted TR sequencing are comparable to the accuracy of PCR sizing analysis of repeats. Overall, we achieved an average genotyping rate of 75% for 110 TR loci with repeat length less than 2kb and GC content greater than 25%. Genotyping rate improved to 91% for the highest-coverage sample, indicating more sequencing could improve genotyping rate.

Targets with low GC sequence content (< 25% GC content) didn’t have sufficient sequence coverage with targeted sequencing. We have previously performed short-read target capture on these regions [12] and observed low sequence coverage in low GC targets. However, both Nanopore and PacBio WGS data didn’t have any bias in sequence coverage in low GC regions. Hence, the lack of sequence coverage in low GC region for targeted sequencing is likely due to the capture protocol. To overcome the issue of low capture efficiency for low GC regions, it is feasible to increase the number of probes in low GC regions during probe design. This will improve sequence enrichment in low GC targets.

We also observed targets greater than 2Kb in length could not be genotyped due to the lack of spanning reads for genotyping analysis. This is primarily due to the limitation in sequence read length observed from the capture process. Streptavidin beads used during the capture process has limitations on the size of the fragments it can bind to, which limits the fragment length attainable with this capture protocol. Although there are longer TRs (greater than 2Kb) in the human genome, more than 99% of the TRs reported in human reference genome (hg38) are less than 2Kb in length [3]. Therefore, our protocol would still be able to successfully genotype most of the TRs in the human genome. TRs greater than 2Kb might need further optimized enrichment protocols.

Our target panel included 8 (out of the 142 targets) STRs with longer expansions (>200 number of copies) and 7 of these targets failed to genotype. However 3 of these had low GC content and one was greater than 4kb in repeat length. The longer expansions which failed to genotype also had low sequence coverage, however due to the low number of targets we couldn’t conclusively identify the cause for failure for these targets.

We used VNTRTyper, an in-house genotyping tool described in Ganesamoorthy et. al. (2018) [12] to determine the repeat number of TRs from long-read sequencing technologies. For comparison, we used Tandem-genotypes [24], recently reported genotyping tool for the detection of TR expansions from long-read sequencing. Both genotyping methods were comparable to PCR sizing analysis and genotyping estimates were comparable between the approaches. However, Tandem-genotypes genotyped less targets than VNTRTyper. The differences is likely due to the different algorithms used between the methods. Both VNTRTyper and Tandem-genotypes uses reads spanning the repeat region. However, for Tandem-genotypes the flanking length used for analysis is depended on the length of the repeat unit, with a maximum of 100bp on both sides of the repeat unit. On the other hand, VNTRTyper uses a default 30bp flanking length for analysis, but it is feasible to change the flanking length. Due to the longer flank length requirement, Tandem-genotypes could have possibly failed to genotype more targets compared to VNTRTyper.

Variations in TRs are a major source of genomic variation between individuals. TRs targeted in this study were initially selected due to the variation observed between case and control samples for obesity analysis [12] and these TRs are variable in the population. We show that approximately 50% of the targeted TRs differ from reported reference copy number. However, the major limitation in this analysis is that the sample size is small and the individuals are related, which introduces a bias in the analysis. Nevertheless, these findings indicate the possibility of variation in TR copy number between individuals and further large-scale studies are required to ascertain the extent of variation.

We demonstrated that the accuracy of genotype estimates between WGS and targeted capture sequencing were comparable to the accuracy of PCR sizing analysis. However, targeted capture enrichment protocols used in this study have amplification steps, which can introduce errors in TR analysis. This could possibly explain the differences in genotype estimates observed between WGS and targeted capture sequencing for some targets.

An amplification free targeted analysis with long-read sequencing is an ideal option for accurate genotyping TRs. Targeted cleavage with Cas9 enzyme followed by Nanopore sequencing [26] or PacBio sequencing [27] has been recently reported as alternative option for enrichment of regions of interest. This method does not have any amplifications and can be adapted for multiple targets in a single assay. However, currently the DNA input requirements are high and sequencing output are low, which currently restricts wide use of this technique for large-scale analysis.

The targeted long-read sequencing approach presented in this study is a cost-effective approach to analyse hundreds of TRs simultaneously. Long-read Nanopore WGS can cost approximately $4000 for 30X coverage of human genome and often with varying coverage across the genome. However, targeted long-read sequencing can be performed for a fraction of cost (less than $300 per sample depending on the multiplexing level) to enrich up to 25MB of genomic sequence of interest. The ability to analyse hundreds of TRs for a fraction of cost allows to explore TRs in large-scale studies.

In summary, we present a targeted approach combined with long-read sequencing to enable cost-effective and accurate approach to genotype TRs using long-read sequencing. Using this method, we have successfully demonstrated the feasibility of targeted capture sequencing of repetitive sequences and genotyping TRs using Nanopore long-read sequencing technology. Our targeted long-read sequencing approach would provide a cost-effective tool for large-scale population analysis of tandem repeats.

## METHODS

### Samples for sequencing

DNA samples of CEPH/UTAH pedigree 1463 were purchased from Coriell Institute for Medical Research (USA). Seven family members from the pedigree used for sequencing analysis were NA12877, NA12878, NA12879, NA12881, NA12882, NA12889 and NA12890.

### Selection of Tandem Repeats and Probe design

The selection of TRs and design of probes were described in Ganesamoorthy *et. al*. (2018) [12]. Briefly, 142 TRs were selected; they range from 112bp to 25236bp in length in the reference human genome (hg19) and the number of repeat units range from 2 to 2300 repeats. TRs used in this study were selected as part of another study to investigate association between TRs and Obesity and these targeted TRs are not disease associated. Agilent SureSelect DNA design (Agilent Technologies) was used to design target probes to capture the targeted regions (including 100bp flanking regions) and regions flanking the TRs (at least 1000bp).

### Nanopore targeted sequencing of TRs

All 7 family members from the CEPH pedigree 1463 were used for Nanopore targeted sequencing analysis (NA12877, NA12878, NA12879, NA12881, NA12882, NA12889 and NA1289). Target sequence capture for Nanopore sequencing was performed using Agilent SureSelect XT HS Target Enrichment System (Agilent Technologies) according to the manufacturer’s instructions with slight modifications. Briefly, 200ng of DNA was fragmented to 3Kb using Covaris Blue miniTUBE (Covaris). Greater than 90% of the targeted TRs are less than 3Kb and SureSelect capture protocol works effectively on fragments less than 4Kb in length; therefore, DNA products were sheared to 3Kb. Fragmented DNA was end repaired, adapter ligated and amplified prior to target capture. Extension time for pre-capture amplification was increased to 4 minutes to allow for the amplification of long fragments and 14 cycle amplification was used. Purified pre-capture PCR products were hybridized to the designed capture probes for 2 hours. Streptavidin beads (Thermo Fisher) were used to pull down the DNA fragments bound to the probes. Finally, captured DNA was amplified with long extension time (4 minutes) using Illumina Index adapters provided in the enrichment kit. Post capture PCR products were purified using 0.8X - 1X AMPure XP beads (Beckman Coulter).

Nanopore sequencing library preparation was performed using 1D Native barcoding genomic DNA (with EXP-NBD103 and SQK-LSK108) (Oxford Nanopore Technologies) protocol according to the manufacturer’s instructions with minor modifications. Briefly, 100ng – 200ng of post capture PCR products were end repaired and incubated at 20°C for 15mins and 65°C for 15mins. End repaired products were ligated with unique native barcodes. Purification steps after end repair and barcode ligation were avoided to minimize the loss of DNA. Barcoded samples were pooled in equimolar concentrations prior to adapter ligation. Adapter ligated samples were purified using 0.4X AMPure XP beads (Beckman Coulter). Samples were split into 2 sequencing groups; NA12877, NA12878, NA12879 and NA12890 – group 1; NA12881, NA12882 and NA12889 – group 2. Sequencing was performed on MinION sequencer (Oxford Nanopore Technologies) using R9.5 flow cell. Both groups were sequenced for 48 hours. Nanopore sequencing data were base called using Albacore (version 2.2.7) and reads were demultiplexed using Albacore (version 2.2.7) based on the barcode sequences.

### Public data used in the study

Nanopore WGS data on CEPH Pedigree 1463 sample NA12878 were obtained from Nanopore WGS consortium [28]. PacBio WGS data on NA12878 sample were downloaded from SRA with accession numbers SRX627421 and SRX638310 [29]

### VNTRTyper

Sequencing reads were mapped to hg19 reference genome using Minimap2 (version 2.13) [30]. For Nanopore sequencing ‘-ax map-ont’ and for PacBio WGS ‘-ax map-pb’ parameters were used. VNTRTyper, our in-house tool described in Ganesamoorthy *et. al.* (2018) [12] was used to genotype TRs from long-read sequencing data. Briefly, VNTRTyper takes advantage of the long-read sequencing to identify the number of repeat units in the TR regions. Firstly, the tool identifies reads that span the repeat region and applies Hidden Markov Models (HMM) to align the repetitive portion of each read to the repeat unit. Then it estimates the multiplicity of the repeat units in a read using a profile HMM.

Recently, we further improved the accuracy of genotyping estimates by clustering the copy number counts from reads to identify the likely genotypes per target. We used Kmeans clustering and the number of clusters are fixed at two clusters for two alleles. A minimum threshold of two supporting reads per genotype was used to assign genotypes. Furthermore, for heterozygous alleles, both alleles should have at least 10% of reads supporting the allele, if not allele with less than 10% of reads was excluded during the analysis. The updated version of VNTRTyper can be accessed from GitHub Japsa release 1.9-3c and can be deployed using script name jsa.tr.longreads. Details of VNTRTyper analysis are previously reported in Ganesamoorthy *et. al.* (2018) [12].

### Tandem-genotypes

We also used another independent method Tandem-genotypes to estimate genotypes from long-read sequencing data. Tandem-genotypes was recently reported for analysis of TR genotypes from long-read sequencing data [24] and it can be utilised for both Nanopore and PacBio sequencing technologies.

Nanopore and PacBio sequencing data were mapped to the hg19 reference genome using LAST v959 [31]. Calculation of repeat length per sequencing read was performed with Tandem-genotypes as reported in Mitsuhashi et. al. (2019) [24]. Copy number changes in reads covering the repeat’s forward and reverse strands were merged and the two alleles with the highest number of supporting reads for each VNTR were extracted. A minimum threshold of two supporting reads per genotype was used to assign genotypes.

### PCR analysis of VNTRs

Ten targeted VNTR regions which are less than 1Kb in repetitive sequence were validated by PCR sizing analysis in this study. These ten targets include various repeat unit length and repeat sequence combinations to assess the accuracy of the genotypes determined from sequencing data. Majority of these targets were tested in our previous study [12] and the results from the previous PCR anlaysis were used for these regions. PCRs were performed using HotStar Taq DNA Polymerase (Qiagen) and PCR conditions were optimized for each PCR target. PCR products were purified and subjected to capillary electrophoresis on an ABI3500xL Genetic Analyzer (Applied BioSystems). Fragment sizes were analyzed using GeneMapper 4.0 (Applied BioSystems).

### Statistical analysis

Linear regression analysis was used to determine correlation between genotype estimates. All plots were generated using GraphPad Prism (version 7.00 for Windows, GraphPad Software, La Jolla California USA).

We investigated the effect of GC content, repeat length, repeat period and repeat copy number on target sequencing depth using a multivariate linear regression model. We used ggplot to visualize the relationship between these factors and sequencing depth across all 7 samples. Thresholds on GC and repeat length were chosen based on this visual analysis. Genotype rate was calculated as the proportion of sample, target pairs which had a predicted genotyped (based on VNTRtyper) amongst all targets which met the GC and repeat length thresholds.

## AVAILABILITY OF DATA AND MATERIALS

The sequencing datasets generated during the study are available in NCBI SRA repository, project number PRJNA422490.

## ACKNOWLEDGEMENTS

This work was supported by the Australian Government National Health and Medical Research Council Project Grant APP1052303 to Prof Lachlan JM Coin. We thank Agilent Technologies for their technical assistance with targeted sequencing experiments.

## AUTHORS’ CONTRIBUTIONS

DG and LC conceived the study. DG, MY and TD performed the experiments. DG, MY, VM, CZ, MDC and LC performed the sequencing and statistical analysis. DG and MY wrote the paper with input from the other authors. All authors read and approved the final manuscript.

## SUPPLEMENTAL DATA

Supplementary.pdf: Includes Supplementary Figures 1 – 5 and Supplementary Table 1-2 Supplementary_Spreadsheets.xls: Includes Tables 1 – 3

