## Supplementary Figure for "High-throughput multiplexed tandem repeat genotyping using targeted long-read sequencing"

**Supplementary Information:**

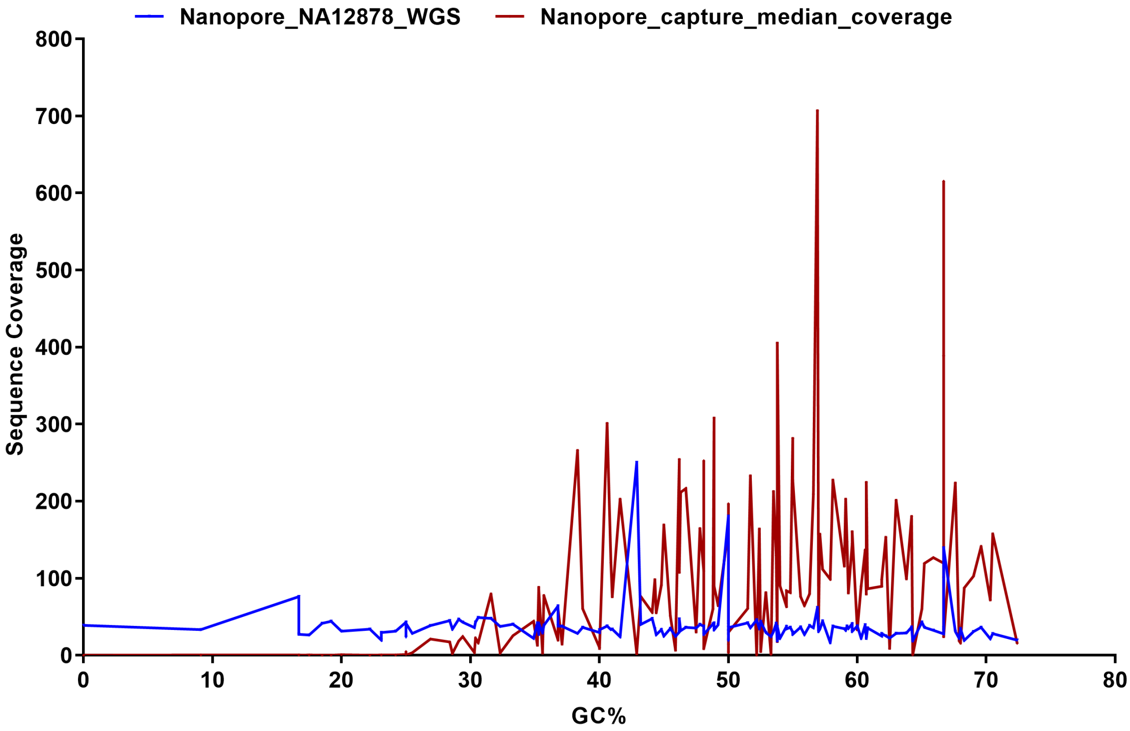


**Supplementary Figure 1**: Sequence coverage distribution on targets with varying GC% for Nanopore sequencing for targeted and whole genome sequencing data. Median sequence coverage of 7 targeted sequencing samples were used.

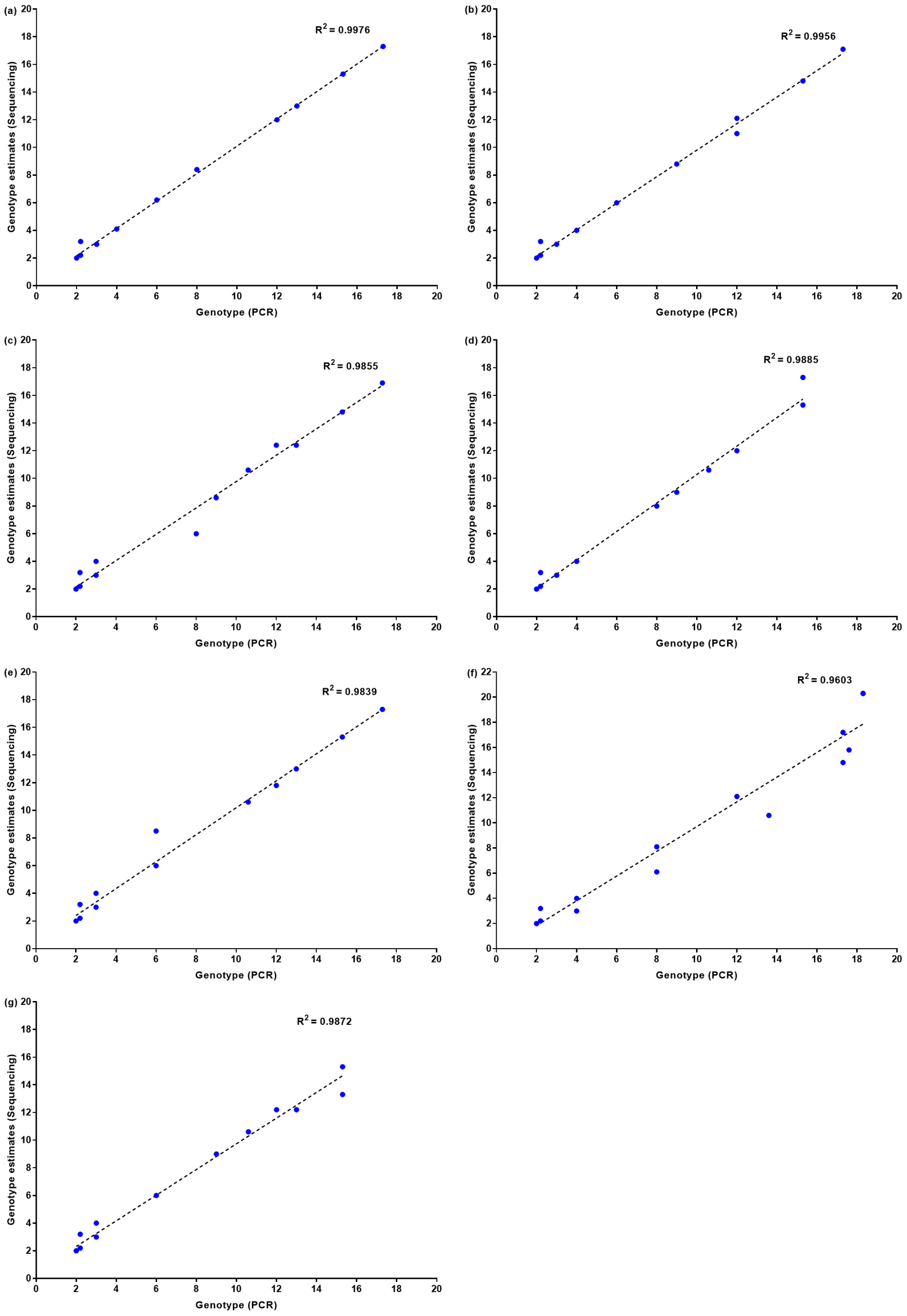


**Supplementary Figure 2**: Correlation of genotype estimates between PCR sizing analysis and genotype estimates determined by VNTRTyper (a) NA128777, (b) NA12878, (c) NA12879, (d) NA12881, (e) NA12882, (f) NA12889 and (g) NA12890.

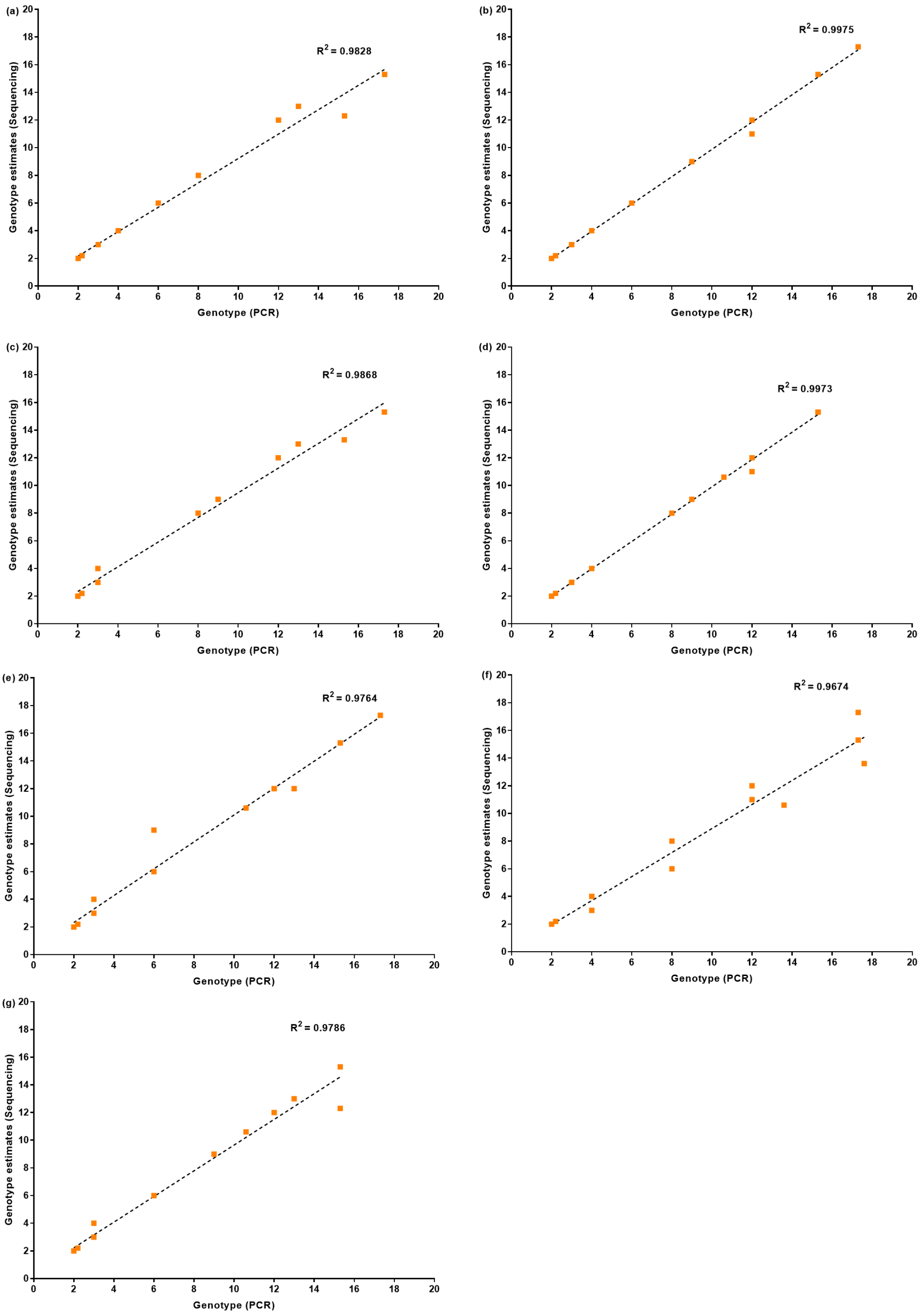


**Supplementary Figure 3**: Correlation of genotype estimates between PCR sizing analysis and genotype estimates determined by Tandem-genotypes (a) NA128777, (b) NA12878, (c) NA12879, (d) NA12881, (e) NA12882, (f) NA12889 and (g) NA12890.

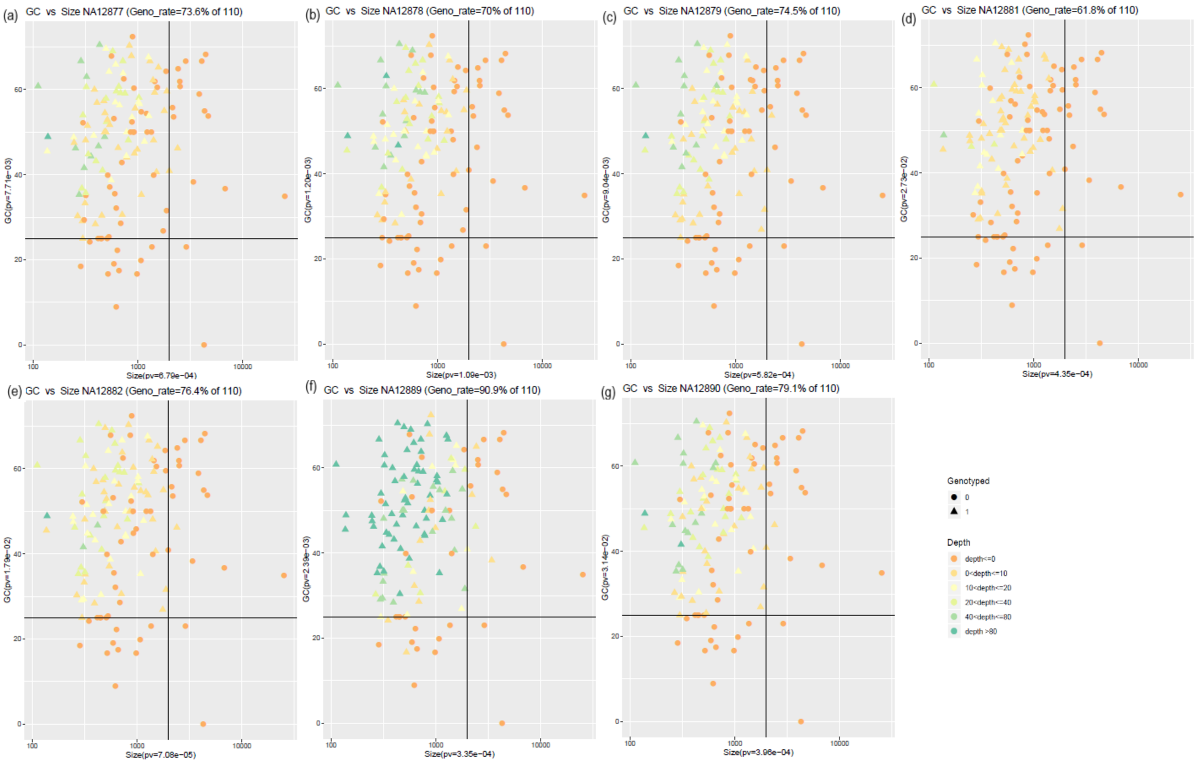


**Supplementary Figure 4:** Genotyping rate with both 25% GC threshold and 2Kb size threshold using VNTRTyper for (a) NA128777, (b) NA12878, (c) NA12879, (d) NA12881, (e) NA12882, (f) NA12889 and (g) NA12890.

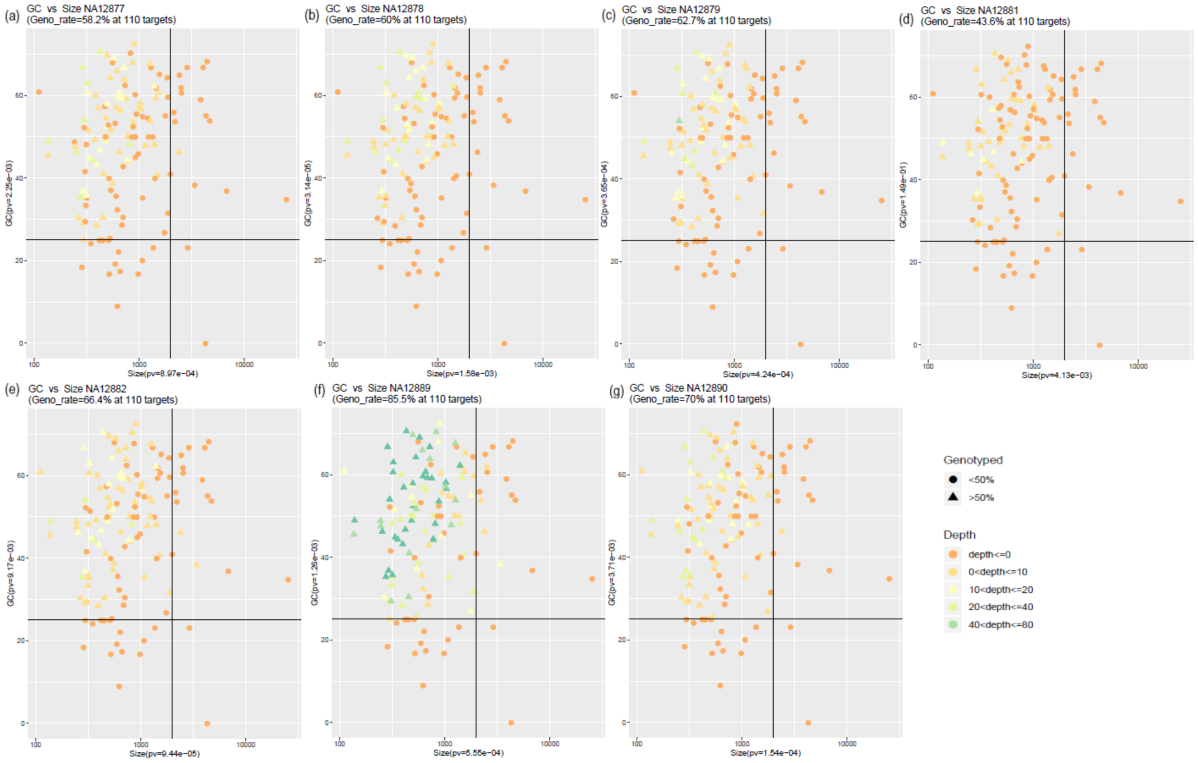


**Supplementary Figure 5:** Genotyping rate with both 25% GC threshold and 2Kb size threshold using Tandem-genotypes for (a) NA128777, (b) NA12878, (c) NA12879, (d) NA12881, (e) NA12882, (f) NA12889 and (g) NA12890.

**Supplementary Table 1**: Targeted Sequencing metrics for Nanopore Capture sequencing of tandem repeats

| Sample | Number of reads | Aligned reads (%) | On-bait bases (%)^ | Off-bait bases (%)^ | Fold enrichment^ | Median Read length (bp) | Max Read length (bp) |
|---|---|---|---|---|---|---|---|
| NA12877 * | 207,976 | 67% | 59% | 41% | 3497 | 568 | 11,207 |
| NA12878 * | 189,050 | 70% | 62% | 38% | 3566 | 646 | 6,781 |
| NA12879* | 209,341 | 70% | 56% | 44% | 3284 | 614 | 24,657 |
| NA12890 * | 206,077 | 76% | 52% | 48% | 2955 | 633 | 8,040 |
| NA12881 # | 44,363 | 77% | 48% | 52% | 2739 | 702 | 5,720 |
| NA12882 # | 106,259 | 77% | 49% | 51% | 2823 | 718 | 7,383 |
| NA12889 # | 1,021,376 | 70% | 45% | 55% | 2573 | 776 | 10,127 |

^ Output from Picard CollectHS Metrics (Version 2.18.29) (On-bait bases - Number of bases that are uniquely mapped to the targeted regions of the genome; Off-bait bases - Number of bases that are uniquely mapped away from the targeted regions of the genome; Fold enrichment - The fold by which the targeted region has been amplified above genomic background)

*Nanopore Multiplexing Group – 1; #Nanopore Multiplexing Group – 2

**Supplementary Table 2**: Genotype estimates on Nanopore targeted capture sequencing using Tandem-Genotypes

| Sample | Method | Genotype of Target* | | | | | | | | Pearson Correlation with PCR |
|---|---|---|---|---|---|---|---|---|---|---|
| | | TR_8 (12.0) | TR_57 (15.6) | TR_86 (2.0) | TR_87 (9.0) | TR_93 (2.0) | TR_109 (15.3) | TR_112 (4.0) | TR_120 (2.2) | |
| **NA12877** | PCR | 12.0/13.0 | 10.6/13.6 | 2.0/2.0 | 6.0/8.0 | 2.0/2.0 | 15.3/17.3 | 3.0/4.0 | 2.2/2.2 | |
| | Nanopore | 12.0/13.0 | ND | 2.0/2.0 | 6.0/8.0 | 2.0/2.0 | 12.3/15.3 | 3.0/4.0 | 2.2/2.2 | 0.9913 |
| **NA12878** | PCR | 12.0/12.0 | 10.6/12.6 | 2.0/2.0 | 6.0/9.0 | 2.0/2.0 | 15.3/17.3 | 3.0/4.0 | 2.2/2.2 | |
| | Nanopore | 11.0/12.0 | ND | 2.0/2.0 | 6.0/9.0 | ND | 15.3/17.3 | 3.0/4.0 | 2.2/2.2 | 0.9988 |
| **NA12879** | PCR | 12.0/13.0 | 10.6/10.6 | 2.0/2.0 | 8.0/9.0 | 2.0/2.0 | 15.3/17.3 | 3.0/3.0 | 2.2/2.2 | |
| | Nanopore | 12.0/13.0 | ND | 2.0/2.0 | 8.0/9.0 | ND | 13.3/15.3 | 3.0/4.0 | 2.2/2.2 | 0.9934 |
| **NA12881** | PCR | 12.0/12.0 | 10.6/10.6 | 2.0/2.0 | 8.0/9.0 | 2.0/2.0 | 15.3/15.3 | 3.0/4.0 | 2.2/2.2 | |
| | Nanopore | 11.0/12.0 | 10.6/10.6 | 2.0/2.0 | 8.0/9.0 | ND | 15.3/15.3 | 3.0/4.0 | 2.2/2.2 | 0.9986 |
| **NA12882** | PCR | 12.0/13.0 | 10.6/10.6 | 2.0/2.0 | 6.0/6.0 | 2.0/2.0 | 15.3/17.3 | 3.0/3.0 | 2.2/2.2 | |
| | Nanopore | 12.0/12.0 | 10.6/10.6 | 2.0/2.0 | 6.0/9.0 | 2.0/2.0 | 15.3/17.3 | 3.0/4.0 | 2.2/2.2 | 0.9881 |
| **NA12889** | PCR | 12.0/12.0 | 13.6/17.6 | 2.0/2.0 | 8.0/8.0 | 2.0/2.0 | 17.3/17.3 | 4.0/4.0 | 2.2/2.2 | |
| | Nanopore | 11.0/12.0 | 10.6/13.6 | 2.0/2.0 | 6.0/8.0 | 2.0/2.0 | 15.3/17.3 | 3.0/4.0 | 2.2/2.2 | 0.9836 |
| **NA12890** | PCR | 12.0/13.0 | 10.6/10.6 | 2.0/2.0 | 6.0/9.0 | 2.0/2.0 | 15.3/15.3 | 3.0/3.0 | 2.2/2.2 | |
| | Nanopore | 12.0/13.0 | 10.6/10.6 | 2.0/2.0 | 6.0/9.0 | 2.0/2.0 | 12.3/15.3 | 3.0/4.0 | 2.2/2.2 | 0.9893 |

* Repeat Number in reference hg19 is provided within brackets for each target

* Repeat numbers that do not agree with PCR results are highlighted in red.

ND – Sufficient data not available for genotype analysis
